# Are new beta-lactamases posing a potential future threat?

**DOI:** 10.1101/2025.02.14.638215

**Authors:** Patrik Mlynarcik, Veronika Zdarska, Milan Kolar

**Affiliations:** Department of Microbiology, Faculty of Medicine and Dentistry, Palacký University Olomouc, Czech Republic

**Author notes:** Corresponding author: Patrik Mlynarcik, Ph.D., Department of Microbiology, Faculty of Medicine and Dentistry, Palacky University Olomouc, Hnevotinska 3, 779 00 Olomouc, Czechia.

**Keywords:** antimicrobial resistance, beta-lactamases, horizontal gene transfer, public health

## Abstract

Antimicrobial resistance, a significant and escalating public health threat, has extended its reach beyond healthcare settings to diverse environments such as urban beaches, wastewater plants, and household pets. A recent study even found bacteria in permafrost with genes encoding beta-lactamases, enzymes that confer resistance to beta-lactam antibiotics. Our previous study identified 2340 potential beta-lactamases across 673 bacterial genera, with only 139 of these enzymes sharing more than 70% amino acid identity with known beta-lactamases. This study underscores the critical and widespread distribution of potential beta-lactamases across different bacterial genera, regions, and hosts. The role of horizontal gene transfer and mobile genetic elements in spreading antibiotic resistance is of utmost importance, highlighting the gravity of the situation and the urgent need for monitoring and research.

## INTRODUCTION

Antimicrobial resistance, a looming and urgent global public health threat, is expected to escalate in severity. Once confined to healthcare settings, antibiotic-resistant bacteria have now proliferated in diverse environments, from washing machines (Schmithausen, Sib et al. 2019), urban beaches (Carney, Labbate et al. 2019), wastewater treatment plants (Parnanen, Narciso-da-Rocha et al. 2019), post-cleaning hospital environments (Gouliouris, Coll et al. 2021), heavy metal environments (Gaeta, Bean et al. 2020), city bird droppings (Zhao, Sun et al. 2020), incinerators and landfills (Li, Wang et al. 2020), dust (Ben Maamar, Glawe et al. 2020), wild dolphins (Schaefer, Bossart et al. 2019), household pets (Hamame, Davoust et al. 2022, Menezes, Moreira da Silva et al. 2022), bedbugs (Herrera, Chaussee et al. 2024), insects and spiders (Hassan, Ijaz et al. 2021), and microplastics (Liu, Liu et al. 2021). This widespread emergence of antibiotic-resistant microbes poses a significant risk to a larger population. A recent study in permafrost (Rigou, Christo-Foroux et al. 2022) even found bacteria with a high proportion of genes encoding beta-lactamases in their genomes, further emphasizing the immediate need for global monitoring and research.

Currently, 346 beta-lactamase types have been described worldwide according to the Beta-Lactamase DataBase (BLDB), a comprehensive and widely recognized repository of beta-lactamase information (http://bldb.eu; last accessed on 10 October 2024). In this context, more than 7280 beta-lactamase genes have been described in cultured bacteria, and new variants are emerging worldwide (Bao, Huang et al. 2022). However, new beta-lactamases are constantly being discovered.

In our previous study (Zdarska, Kolar et al. 2024), silico analysis of over 850 bacterial genomes from clinical and environmental isolates revealed more than 2340 potential beta-lactamases in over 673 bacterial genera. In the present study, we focused more on candidate beta-lactamases with 70% or greater amino acid sequence identity with known beta-lactamases.

## METHODS

For the analysis, 139 bacterial genomes and gene sequences encoding putative beta-lactamases were downloaded from the National Center for Biotechnology Information (NCBI) and examined. These sequences, annotated using the NCBI prokaryotic genome annotation pipeline (Tatusova, DiCuccio et al. 2016), were filtered using the “find annotations” feature in Geneious Prime 2024.0.7 (Kearse, Moir et al. 2012).

This study only considered sequences with 70.0-92.3% amino acid identity with known beta-lactamases. The amino acid identity threshold of 92.3% was used to identify potential beta-lactamases as it indicates significant similarity to known beta-lactamases, suggesting a high likelihood of functional similarity. Amino acid matches were determined using the BLDB BLAST function (Naas, Oueslati et al. 2017).

The flanking regions of beta-lactamases were analyzed using Geneious Prime. The NCBI Protein database and Geneious data were also utilized to obtain information on the country of origin, host, and source of isolation of the bacterial isolates.

## RESULTS AND DISCUSSION

One hundred thirty-nine beta-lactamases in this study exhibited ≥ 70% amino acid identity with known beta-lactamases. The distribution of potential beta-lactamases across various regions and hosts is shown in Figure 1. In total, we identified 109 bacterial genera and one *Candidatus Ornithobacterium* species (see Table S1), comprising the following bacterial classes (*Actinomycetes, Alphaproteobacteria, Bacilli, Bacteroidia, Betaproteobacteria, Cyanophyceae, Cytophagia, Deltaproteobacteria, Epsilonproteobacteria, Flavobacteriia, Gammaproteobacteria, Chitinophagia, Chlorobiia*, and *Sphingobacteriia*), in which the following novel potential beta-lactamases of group A (53 times), group B (27 times), group C (12 times) and group D (47 times) were detected.

**Figure 1.**
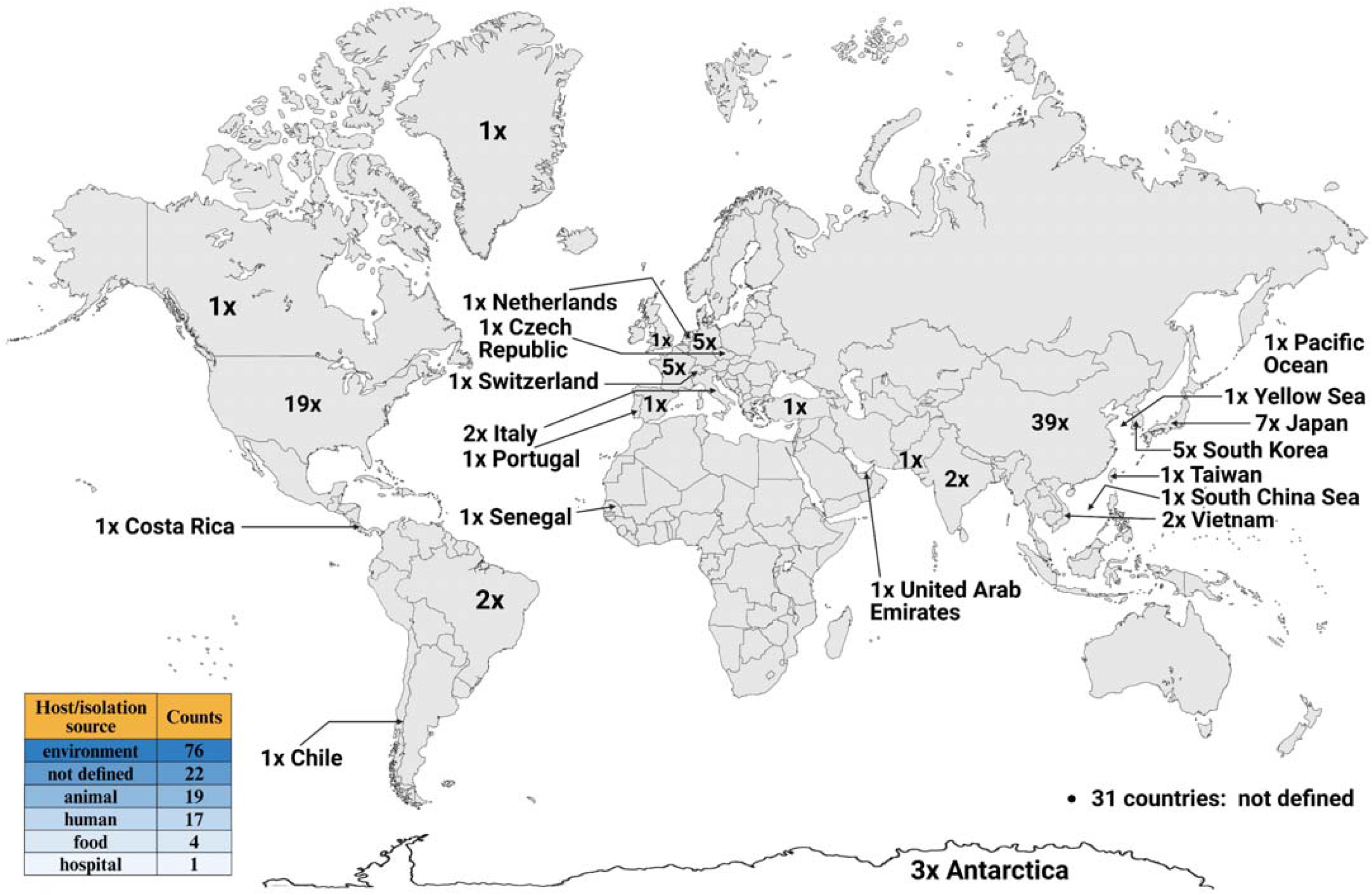
Global distribution of potential beta-lactamases in different regions and hosts. Created in BioRender.com.

The highest numbers of potential beta-lactamases were found in the genera *Chryseobacterium, Enterobacter, Aeromonas* and *Chitinophaga*, in which six, four and three enzymes (twice), respectively, were detected. For example, two potential class A beta-lactamases (83.2% and 80.5% amino acid identity with CGA-1), three class B beta-lactamases (87.9% identity with ESP-1; 71.6% and 70.7% identity with GOB-like enzymes), and one class D beta-lactamase (89% identity with OXA-209) were detected in the genus *Chryseobacterium*. Further, in the genus *Enterobacter*, there were two potential class A beta-lactamases (89.1% and 82.5% amino acid identity with LAP-1 and PLA-6, respectively) and two class C beta-lactamases (87.1% and 85.6% identity with ACT-96 and CDA-1, respectively). In the case of the genus *Aeromonas*, there was one potential beta-lactamase of class A (70.6% amino acid identity with BlaP-2), class C (86.4% identity with PAC-1), and class D (86.2% identity with OXA-12).

Identification of mobile genetic elements revealed the presence of genes for transposases, integrases/recombinases, and mobilization proteins in 14 cases (Table 1), with a higher abundance of transposases, which may lead to overproduction of beta-lactamases and facilitate gene transfer, spreading beta-lactamase genes. In addition, two potential beta-lactamases were detected nearby in some cases (Figure 2A/B).

**Table 1.**
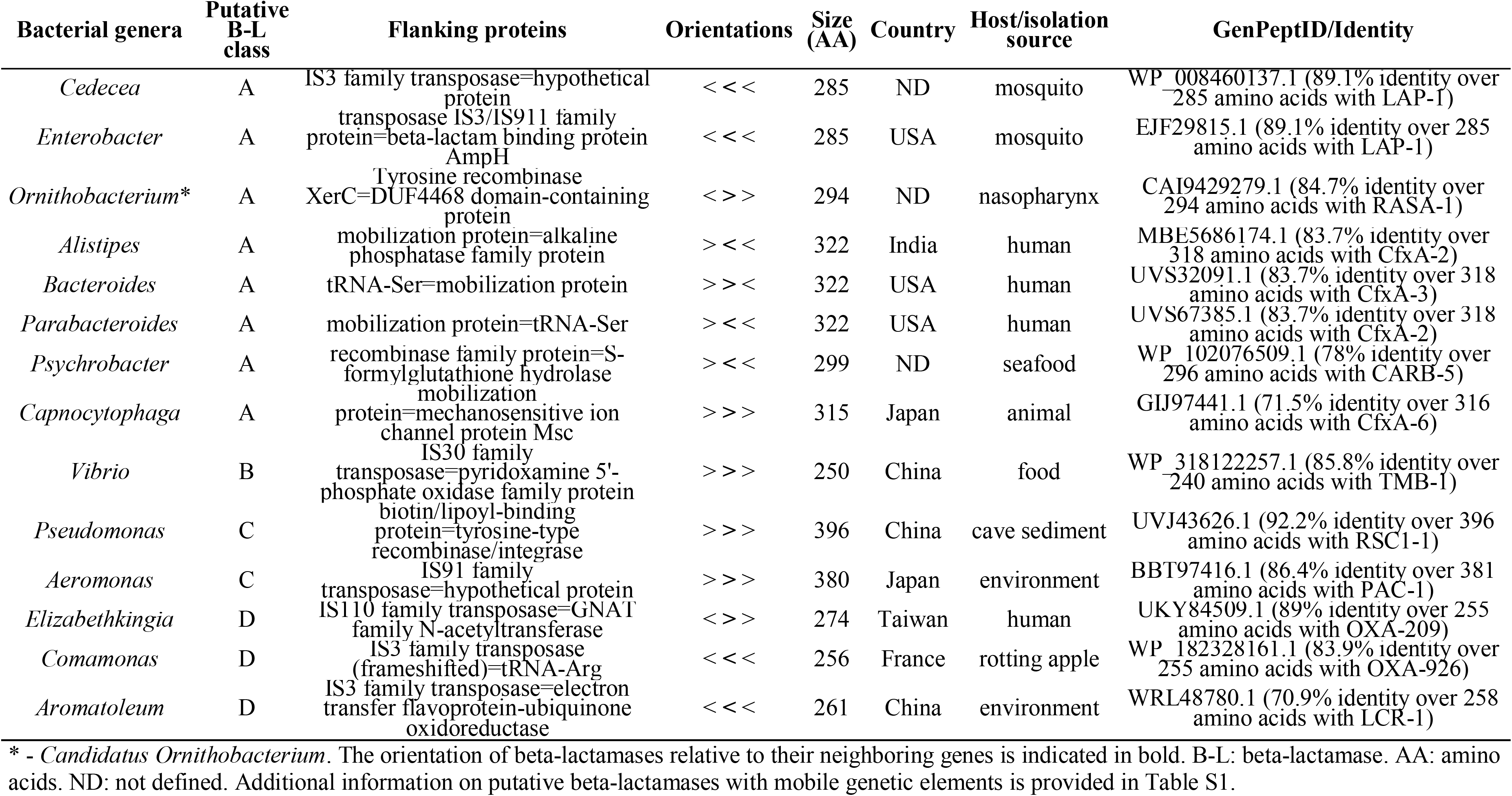
Genetic neighborhood of selected putative beta-lactamases with mobile genetic elements.

**Figure 2.**
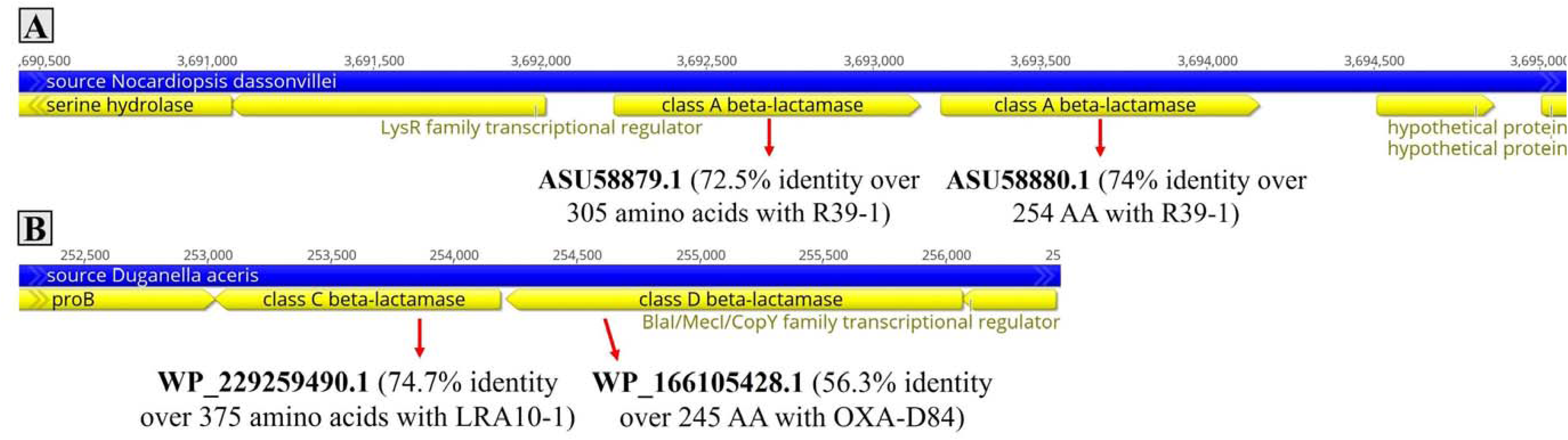
Detection of potential beta-lactamases in *Nocardiopsis dassonvillei* (A) and *Duganella aceris* (B). Arrows indicate the direction of transcription. The length of the arrows is proportional to the size of the genes.

The study analyzed the presence of potential beta-lactamases in bacterial isolates from various countries and demonstrated a notable distribution of these enzymes across different regions. China had the highest number of potential beta-lactamases, with 39 cases detected. In addition, there were 31 cases where the source country was not specified. The USA reported 19 cases, Japan had seven cases, and France, Germany, and South Korea each had five cases. Additionally, we explored the presence of potential beta-lactamases across different isolation environments. The majority were detected in environmental samples, accounting for 76 cases, followed by 22 cases where the origin was not determined. Furthermore, 19 cases were identified in animal and 17 in human hosts.

Our detailed study of potential beta-lactamases binding to mobile genetic elements resulted in a significant discovery. Specifically, we identified two potential class A beta-lactamases with identical amino acid sequences in *Enterobacter* sp. (AKXM01000000) and *Cedecea* sp. (NZ_JBBCPU010000048), both of which were found in mosquitoes and had an insertion sequence located nearby. This finding is crucial as it highlights potential sources of antibiotic resistance, with one case originating from the USA and the other from an unspecified location.

An insertion sequence, a short DNA segment that acts as a simple transposable element, was also found around potential class C beta-lactamase described in *Aeromonas veronii* (AP022282) isolated from a wastewater treatment plant effluent in Japan. This insertion sequence has also been detected in other bacterial species of this genus, such as *Aeromonas caviae* (AP022242, AP019196), all isolated from the same source and country.

The most worrying situation seems to be related to potential class A beta-lactamases, which show 99.9% nucleotide and 99.7-100% amino acid identity (change in one amino acid) with each other and have been detected in *Bacteroides faecis* (CP103274), *Parabacteroides distasonis* (CP103286) and *Alistipes* sp. (RKBL01000025), and these enzymes were located near the mobilization protein. More detailed *in silico* analysis indicated their occurrence in other species of the above genera, such as *Bacteroides ovatus* (CP103071), *Bacteroides fragilis* (CP036539, CP119600), *Bacteroides salyersiae* (CP072243), *Bacteroides* sp. (CP040630), *Parabacteroides distasonis* (CP072231), and *Parabacteroides goldsteinii* (CP081906). In addition, these enzymes have also been detected in other bacterial species, such as *Phocaeicola vulgatus* (CP096965) and *Prevotella* sp. (AP035786). These enzymes originated from humans and were found in India, the USA, Canada, Denmark, and the Netherlands, and from horses (*Equus caballus*) in Japan. This fact illustrates the seriousness of the situation, as the presence of beta-lactamases with a mobilization protein has been detected in many genera and species, indicating their widespread distribution and potential for the spread of antibiotic resistance.

Our study highlights the significant distribution of potential beta-lactamases across different bacterial genera, regions, and hosts. The presence of these enzymes, with exact amino acid identities with known beta-lactamases, in genera such as *Cedecea, Enterobacter, Alistipes, Bacteroides*, and *Parabacteroides*, suggests that horizontal gene transfer may play a role in spreading antibiotic resistance. This is supported by the identification of mobile genetic elements surrounding these genes, which may facilitate their transfer. China had the highest number of potential beta-lactamases, with the majority detected in environmental samples. The detection of these enzymes in various environmental and clinical settings underscores the need for comprehensive monitoring and international surveillance. To combat antibiotic resistance, global cooperation is needed. Mobile genetic elements, such as transposases, highlight the potential for these resistance genes to spread. This is crucial for predicting treatment failures and understanding the spread of resistance within and across bacterial species. The presence of beta-lactamases in diverse environments suggests that natural habitats may serve as reservoirs for these resistance genes, potentially allowing them to be transferred to clinical settings. Environmental strains provide a significant reservoir of new resistance genes, including carbapenemases, which can be disseminated through the food chain, accelerating global antibiotic resistance. Human activities facilitate the spread of resistance from various reservoirs such as washing machines and urban beaches, contributing to the complexity of combating antibiotic resistance. This broad distribution underscores the severity of the issue and the need for further research to understand and tackle antimicrobial resistance. This is also why we are convinced that we will soon witness the discovery of numerous new types of beta-lactamases. Many of these previously undescribed beta-lactamases may also play a role in the resulting resistance to various beta-lactam antibiotics. This emphasizes the urgent need for continued surveillance and research to combat antibiotic resistance and protect public health. Addressing this challenge requires the integration of clinical and environmental perspectives to effectively monitor and mitigate the impact of beta-lactamases on global health.

## SUPPLEMENTARY DATA

Table S1: Summary of putative beta-lactamases in various bacterial classes.

## DECLARATION OF GENERATIVE AI AND AI-ASSISTED TECHNOLOGIES IN THE WRITING PROCESS

During the preparation of this work, the author(s) used ChatGPT to improve readability and language. After using this tool/service, the author(s) reviewed and edited the content as needed and take(s) full responsibility for the publication’s content.

## ACKNOWLEDGEMENTS

This work was supported by the project National Institute of Virology and Bacteriology (Programme EXCELES, ID Project No. LX22NPO5103), funded by the European Union – Next Generation EU, and the Internal Grant Agency of Palacký University (project IGA_LF_2025_022).

